# Tumor necrosis factor-α modulates GABAergic and Dopaminergic neurons in the ventral periaqueductal gray of female mice

**DOI:** 10.1101/2021.06.02.446764

**Authors:** Dipanwita Pati, Thomas L. Kash

**Affiliations:** Bowles Center for Alcohol Studies, University of North Carolina at Chapel Hill, Thurston Bowles Building 104 Manning Drive, Chapel Hill, NC, 27599, USA; Department of Pharmacology, University of North Carolina School of Medicine, Chapel Hill, NC, 2751, USA

**Keywords:** PAG, TNF-α, Excitability, GABA, Dopamine, Glutamate, synaptic transmission

## Abstract

Neuroimmune signaling is increasingly identified as a critical component of various illnesses, including chronic pain, substance use disorder, and depression. However, the underlying neural mechanisms remain unclear. Proinflammatory cytokines, such as tumor necrosis factor-α (TNF-α), may play a key role by modulating synaptic function and long-term plasticity. The midbrain structure periaqueductal gray (PAG) plays a well-established role in pain processing, and while TNF-α inhibitors have emerged as a potential therapeutic strategy for pain-related disorders, the impact of TNF-α on PAG neuronal activity has not been thoroughly characterized. Recent studies have identified subpopulations of ventral PAG (vPAG) with opposing effects on nociception, with DA neurons driving pain relief in contrast to GABA neurons. Therefore, we used ex vivo slice physiology to examine the effects of TNF-α on neuronal activity of both subpopulations. We selectively targeted GABA and dopamine neurons using a vGAT-reporter and a TH-eGFP reporter mouse line, respectively. Following exposure to TNF-α, the intrinsic properties of GABA neurons were altered, resulting in increased excitability along with a reduction in glutamatergic synaptic drive. In DA neurons, TNF-α exposure resulted in a robust decrease in excitability along with a modest reduction in glutamatergic synaptic transmission. Furthermore, the effect of TNF-α was specific to excitatory transmission onto DA neurons as inhibitory transmission was unaltered. Collectively, these data suggest that TNF-α differentially affects the basal synaptic properties of GABA and DA neurons and enhances our understanding of how TNF-α mediated signaling modulates vPAG function.

**New & Noteworthy:** The present study describes the effects of tumor necrosis factor-α (TNF-α) on two distinct subpopulations of neurons in the ventral periaqueductal gray (vPAG). We show that TNF-α alters both neuronal excitability and glutamatergic synaptic transmission on GABA neurons and dopamine neurons within the vPAG. This provides critical new information on the role of TNF-α in the potential modulation of pain since activation of vPAG GABA neurons drives nociception, whereas activation of DA neurons drives analgesia.

## Introduction

The midbrain periaqueductal gray area (PAG) is an evolutionarily conserved region that regulates a wide range of complex behaviors, including pain, arousal, and fight-or-flight behaviors (Bandler and Keay 1996; Basbaum and Fields 1978; Graeff et al. 1993). The ventral column of the PAG (vPAG) is a major site of endogenous opioid-induced anti-nociception mediated through its descending projections via the rostral ventromedial medulla (RVM) (Behbehani 1995; Fields 2004; Moreau and Fields 1986). The vPAG comprises heterogeneous subpopulations of neurons that modulate divergent behaviors. For example, it has been shown that vPAG glutamate neurons promote anti-nociception and escape behaviors in contrast to vPAG GABA neurons (Samineni et al. 2017; Tovote et al. 2016; Zhu et al. 2019). Another understudied subpopulation of neurons within the vPAG/dorsal raphe are dopamine (DA) neurons. vPAG DA neurons are a subset of glutamate neurons that co-release glutamate and have been implicated in anti-nociceptive effects (Flores et al. 2004; Li et al. 2016; Suckow et al. 2013; Taylor et al. 2019), fear learning (Groessl et al. 2018; Tovote et al. 2016), arousal (Cho et al. 2017; Porter-Stransky et al. 2019)and incentivized salience (Lin et al. 2020). Also, recent work from our lab (Yu et al. 2021) has shown that either chemogenetic activation of vPAG DA cells or optogenetic stimulation of vPAG terminals in the extended amygdala reduces nociceptive sensitivity during naïve and inflammatory pain states in males but not in females.

Recent evidence has implicated glial-mediated neuroinflammatory response in the pathogenesis of chronic pain and morphine tolerance. Repeated administration of morphine results in upregulated expression of proinflammatory cytokines (Raghavendra et al. 2002; Song and Zhao 2001). In rats, withdrawal from chronic morphine results in upregulation of a proinflammatory cytokine, tumor necrosis factor-alpha (TNF-α) in the ventrolateral PAG, and microinjection of recombinant TNF-α resulted in morphine-withdrawal like behavioral signs (Hao et al. 2011). Furthermore, there’s evidence demonstrating that PAG TNF-α signaling decreases the efficacy of opioids by promoting neuroinflammation and disrupting glutamate homeostasis (Eidson et al. 2017; Eidson and Murphy 2013). Work from the same group also suggests increased activation of PAG microglia in females contributes to sex differences in morphine analgesia (Doyle et al. 2017). These results support the hypothesis that proinflammatory cytokine signaling in the PAG may play an important role in the sex-dependent modulation of pain through opioid signaling (for an extensive review, see (Averitt et al. 2019).

While there have been studies looking at how TNF-α can regulate synaptic plasticity and neuronal excitability in different brain regions via modulation of AMPA receptor and glutamate homeostasis (Lewitus et al. 2014; Shim et al. 2018; Stellwagen et al. 2005a; Stellwagen and Malenka 2006), there has been no direct evidence showing TNF-α-mediated changes in excitability and glutamatergic transmission in the vPAG. Given that pain affects women disproportionately (Mogil 2012), more insight into the synaptic effects of TNF-α in female subjects is warranted. To address this disparity, in the present study, we utilized whole-cell patch clamping to investigate the role of TNF-α in the vPAG of female mice. We provide a novel characterization of the synaptic effects of TNF-α signaling on two specific subpopulations of vPAG neurons: DA and GABA neurons that play critical roles in eliciting a diverse set of nociceptive and aversive behaviors.

## Materials and Methods

### Mice

All experiments were performed on adult female mice (2-5 months) in accordance with the NIH guidelines for animal research and with the approval of the Institutional Animal Care and Use Committee at the University of North Carolina at Chapel Hill. Animals were group-housed in a ventilated and temperature-controlled vivarium on a standard 12-hour cycle (lights on at 07:00) with *ad libitum* access to food and water. To visualize vGAT-expressing neurons, vGAT-ires-Cre mice were crossed with a Cre-inducible L10-GFP reporter line(Krashes et al. 2014) to produce vGAT-L10A-GFP mice. Female TH-eGFP mice on a Swiss Webster background were used for visualizing dopamine neurons (Li et al. 2016). In the TH-eGFP mouse line, the genome was modified to contain multiple copies of a modified BAC in which an eGFP reporter gene was inserted immediately upstream of the coding sequence of the gene for tyrosine hydroxylase (TH). Data presented here were obtained from transgenic mice maintained in-house.

### Slice electrophysiology

For *ex vivo* slice physiology, mice were anesthetized with isoflurane and were rapidly decapitated. 250 μm thick coronal sections through the PAG were prepared as previously described (Pati et al. 2019). Briefly, brains were quickly extracted, and slices were made using a Leica VT 1200s vibratome (Leica Biosystems, IL, USA) in ice-cold, oxygenated sucrose solution containing in mM: 194 sucrose, 20 NaCl, 4.4 KCl, 1 MgCl_2_, 1.2 NaH_2_PO_4_, 10 glucose and 26 NaHCO_3_ saturated with 95 % O_2_/5 % CO_2_. Slices were incubated for at least 30 minutes in normal artificial cerebral spinal fluid (ACSF) maintained at 32-35°C that contained in mM: 124 NaCl, 4.0 KCl, 1 NaH_2_PO_4_, 1.2 MgSO_4_, 10 D-glucose, 2 CaCl_2_, and 26 NaHCO_3_, saturated with 95% O_2_/5 % CO_2_ before transferring to a submerged recording chamber (Warner Instruments, CT, USA) for experimental use. For whole-cell recordings, slices were continuously perfused at a rate of 2.0-3.0 ml/min with oxygenated ACSF maintained at 30±2°C.

Neurons were identified using infrared differential interference contrast on a Scientifica Slicescope II (East Sussex, UK). Fluorescent cells were visualized using a 470 nm LED. Whole-cell patch clamp recordings were performed using micropipettes pulled from a borosilicate glass capillary tube using a Flaming/Brown electrode puller (Sutter P-97; Sutter Instruments, Novato, California). Electrode tip resistance was between 3 and 6 MΩ. All signals were acquired using an Axon Multiclamp 700B (Molecular Devices, Sunnyvale, CA). Data were sampled at 10 kHz, low pass filtered at 3 kHz. Access resistance was continuously monitored, and changes greater than 20% from the initial value were excluded from data analyses. 2-4 cells were recorded from each animal per set of experiments.

Excitability experiments were performed in current-clamp mode using a potassium gluconate-based intracellular solution (in mM: 135 K-gluconate, 5 NaCl, 2 MgCl_2_, 10 HEPES, 0.6 EGTA, 4 Na_2_ATP, 0.4 Na_2_GTP, pH 7.3, 285–295 mOsm). Input resistance was measured immediately after breaking into the cell. Following stabilization, current was injected to hold cells at a common membrane potential of -70 mV to account for inter-cell variability. Changes in excitability were evaluated by measuring rheobase (minimum current required to elicit an action potential), action potential (AP) threshold, and the frequency of action potentials fired at increasing 20 pA current steps (0 to 200 pA). Parameters related to AP kinetics, which included AP height, AP duration at half-maximal height (AP half-width), time to fire an AP (AP latency), and afterhyperpolarization amplitude (AHP), were calculated from the first action potential fired during the F-I plot (Bean 2007). AP height was calculated as the difference between AP peak and AP threshold voltage. AP latency was defined as the duration to the first AP following a depolarizing current step of fixed amplitude. AP half-width was measured as the AP duration at the membrane voltage halfway between AP threshold and AP peak. AHP was calculated as the difference from the action potential threshold to the minimum voltage in the 25 ms following the action potential. AP depolarization and repolarization rates were calculated from the first derivative of its voltage recording with respect to time.

For assessment of spontaneous synaptic activity, two different intracellular solutions were used. Spontaneous excitatory postsynaptic currents (sEPSCs) were assessed in voltage clamp using a potassium gluconate-based internal (in mM: 135 K-gluconate, 5 NaCl, 2 MgCl_2_, 10 HEPES, 0.6 EGTA, 4 Na_2_ATP, 0.4 Na_2_GTP, pH 7.3, 285–290 mOsm). Cells were voltage-clamped at -80 mV in the presence of 25 μM picrotoxin (GABA-A receptor antagonist) to pharmacologically isolate EPSCs. Spontaneous inhibitory postsynaptic currents (sIPSCs) were pharmacologically isolated by adding kynurenic acid (3mM) to the ACSF to block AMPA and NMDA receptor-dependent postsynaptic currents. Cells were clamped at -70 mV and recorded using a potassium-chloride gluconate-based intracellular solution (in mM: 80 KCl, 70 K-gluconate, 10 HEPES, 1 EGTA, 4 Na_2_ATP, 0.4 Na_2_GTP, pH 7.2, 285–290 mOsm with 1mg/ml QX-314-bromide). Cumulative probability plots were constructed to compare the effects of TNF-α on the distribution of amplitude and inter-event intervals (IEI) from sEPSCs (McQuiston and Colmers 1996). IEI histograms were binned at 20 ms, and amplitude histograms were binned in 5 pA intervals. Synaptic drive was defined as the product of sEPSC frequency and amplitude.

### TNF-α incubation

Slices were preincubated in either ACSF alone or ACSF + TNF-α (100 ng/ml) for 1 h before transferring to a submerged recording chamber, similar to (Lewitus et al. 2014; Shim et al. 2018). For whole-cell recordings, slices were continuously perfused with either oxygenated ACSF (Control group) or ACSF + TNF-α (TNF-α group).

### Drugs

All chemicals used for slice electrophysiology were obtained from either Tocris Bioscience (Minneapolis, USA) or Abcam (Cambridge, UK). Recombinant human TNF-α protein was purchased from R&D Systems (Minneapolis, USA) and stored at -20°C.

### Data and statistical analysis

Differences in various electrophysiological measures were analyzed in Clampfit 10.7 (Molecular Devices, Sunnyvale, CA) and compared between the Control and TNF-α groups. Frequency, amplitude, and kinetics of E/IPSCs were analyzed and visually confirmed using Clampfit 10.7. For comparisons between two groups, P-values were calculated using a standard unpaired t-test. If the condition of equal variances was not met, Welch’s correction was used. Cumulative probability plots were constructed to compare the effects of TNF-α on the distribution of amplitude and inter-event intervals from sEPSCs. Amplitude histograms were binned in 5 pA intervals. Comparisons between the groups in the distribution of sEPSCs amplitudes and inter-event intervals were made by performing the Kolmogorov-Smirnov test. Repeated measures ANOVAs (treatment X current injection) were used to assess between-group differences in the spike frequency fired across a range of current steps. All data are expressed as mean ± SEM. P-values ≤ 0.05 were considered significant. All statistical analysis was performed using GraphPad Prism v.8 (La Jolla, CA, USA).

## Results

### TNF-α increases the excitability of vPAG GABA neurons

To determine the effects of TNF-α on the excitability of vPAG GABA neurons, we conducted whole-cell recordings from vGAT neurons in vGAT-L10a reporter mice (Fig. 1A; 4-5 mice per group). Intrinsic properties of vGAT neurons were calculated from data generated in voltage clamp using steps from -70 to -80 mV. The average capacitance was generally < 30 pF and did not differ between the two groups (Fig.1B; n=12 cells in Control group; n=8 cells in the TNF-α group; p=0.2301). We found that the average input resistance in the TNF-α group Control group was slightly higher than the Control group but was not statistically different (Fig.1C; p=0.0808). Next, we performed voltage clamp experiments to evaluate the relationship between holding current and command potential between -70 and -120 mV with 10 mV current steps (Fig.1D). A two-way repeated-measures ANOVA revealed a significant group x voltage interaction [F(5,90)=3.316; p=0.0085], along with a main effect of voltage [F(1.068, 19.22)=57.43; p=<0.0001] and a strong trend toward main effect of group [F(1, 18)=4.370; p=0.0510]. To further investigate the effect of TNF-α on neuronal excitability, we assessed excitability parameters in current-clamp mode. All measures of excitability were taken at -70 mV to normalize for inter-cell variability in RMP. The action potential threshold and the amount of current required to fire an action potential (rheobase) were assessed through a ramp protocol of 120 pA/1s. TNF-α significantly reduced the rheobase compared to the Control group (Fig.1 E-F; p=0.0296; unpaired t-test with Welch’s correction) with no change in the action potential threshold (Fig. 1G; p = 0.640). Next, we measured the action potential frequency across a range of current steps (0-200 pA for 500 ms, at an increment of 20 pA). There was no significant interaction between the two groups as revealed by repeated-measures two-way ANOVA (Fig. 1H-I; [F(10,180)=0.1885, p=0.9969] for group x current interaction; [F(1,18)=0.3953, p=0.5374] for main effect of group; [F(1.848,33.26)= 44.49, p<0.0001] for main effect of current). Additionally, TNF-α failed to alter action potential kinetics of vGAT neurons. Specifically, both groups were similar with respect to AP latency (Fig.1J; p=0.844), AP height (Fig.1K; p=0.1528), AP half-width (Fig.1L; p=0.2032), AHP (Fig.1M; p=0.8328), depolarization rate (Fig.1N; p=0.5875) and repolarization rate (Fig.1O; p=0.1993). Together these results suggest that TNF-α incubation results in alterations of intrinsic properties of vGAT neurons, potentially increasing the excitability of these neurons.

**Figure 1.**
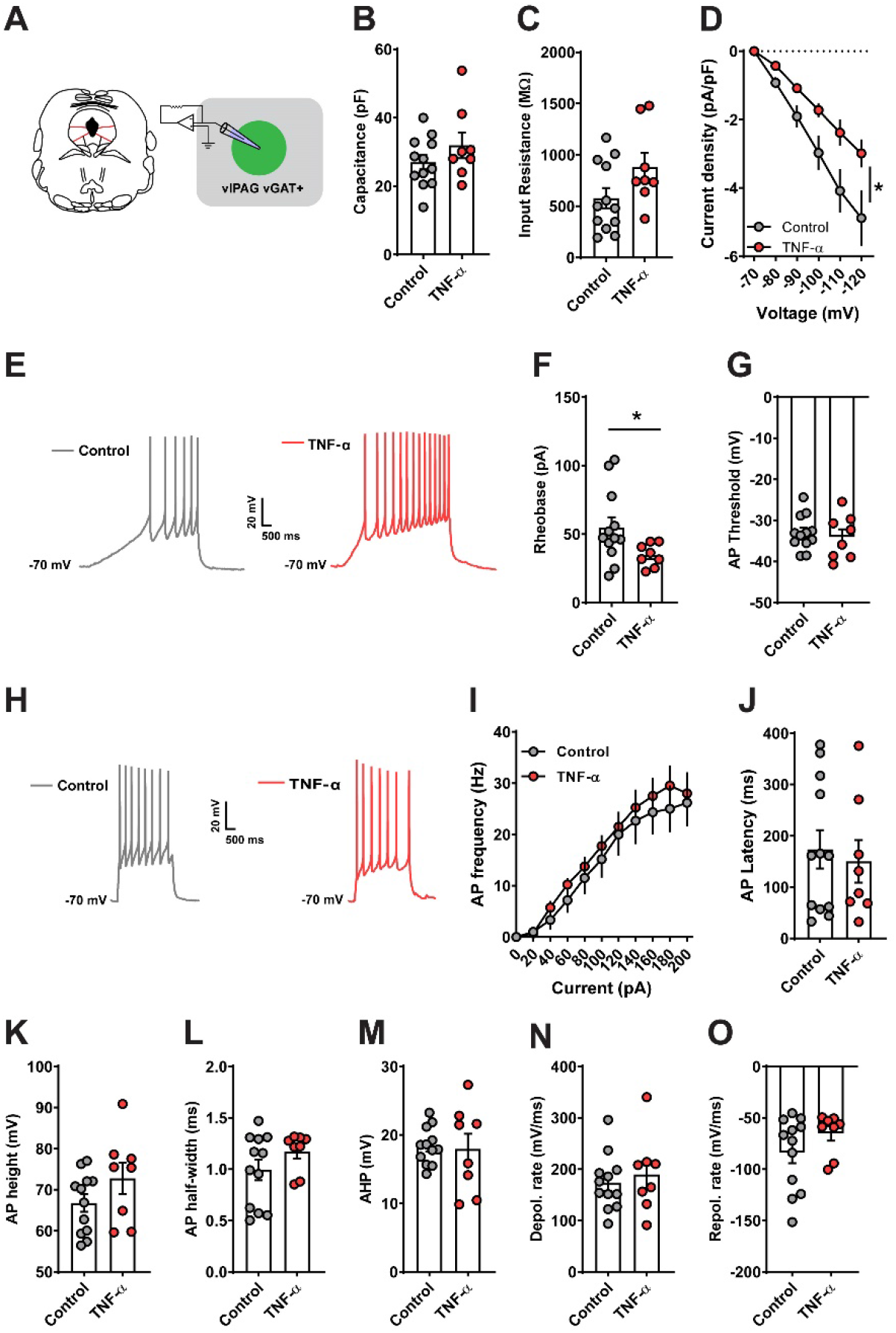
TNF-α incubation increases excitability of vPAG GABA neurons. **(A)** Schematics for whole-cell recordings from vPAG vGAT neurons (n=4-5 mice in each group) following pre-incubation of slices in TNF-α (100 ng/ml). **(B-C)** There was no change in cell capacitance or input resistance between the two groups. **(D)** TNF-α resulted in a reduction in current density in response of varying changes in voltage. **(E)** Representative data obtained from vGAT neurons in in response to a 120 pA/s current ramp while injecting a constant current to hold the cells at -70 mV. The minimum current required to fire an action potential (rheobase) was reduced following pre-incubation in TNF-α **(F)** without any changes **(G)** in AP threshold. **(H)** Representative traces of action potentials fired in response to a step protocol of increased current steps of 20 pA/500 ms while holding the cells at -70 mV. **(I)** There was no significant interaction between the frequency of APs in response to a graded current injection. TNF-α did not alter action potential kinetics as measured by latency **(J)**, average AP height **(K)**, AP half-width **(L)**, AHP **(M)**, depolarization rate **(N)**, and repolarization rate **(O)**. Data expressed as Mean ± SEM.*p<0.05.

### TNF-α decreases neuronal excitability of vPAG Dopamine neurons

We next asked whether TNF-α might also modulate the excitability of vPAG DA neurons. vPAG DA neurons were identified using TH-eGFP reporter mice (Fig. 2A; 4-6 mice per group). There were no significant differences between the two groups with regard to capacitance (Fig. 2B; n=10 cells per group; p=0.1384; unpaired t-test with Welch’s correction), input resistance (Fig. 2C; p=0.2339) and current-voltage relationship (Fig. 2D; two-way repeated-measures ANOVA revealed a significant effect of voltage [F(1.136, 20.44)=89.17; p=<0.0001] but no main effect of group [F(1,18)=0.0018; p=0.9668] or interaction [F(5,90)=0.017; p=0.9999]). Interestingly, we observed a significant increase in rheobase (Fig. 2E-F; p=0.0291) and a non-significant trend toward an increase in AP threshold (Fig. 2G; p=0.0759; unpaired t-test with Welch’s correction). Further, we found a significant reduction in action potential frequency (Fig. 2H-I). A two-way repeated-measures ANOVA revealed a significant interaction between the two groups and the AP frequency [F(9,162)=2.217; p=0.0234], main effect of group [F(1,18)=5.675; p=0.0284] and main effect of current [F(1.310, 23.58)=23.56; p=<0.0001]. Similar to vGAT neurons, there was no impact of TNF-α on AP kinetics as measured by AP latency (Fig.2J; p=0.7193), AP height (Fig.2K; p=0.1714), AP half-width (Fig.2L; p=0.1805), AHP (Fig.2M; p=0.4687), depolarization rate (Fig.2N; p=0.2633) and repolarization rate (Fig.2O; p=0.4493). These results collectively show that TNF-α reduces the excitability of vPAG DA neurons suggesting differential modulation of two distinct subpopulations in the vPAG.

**Figure 2.**
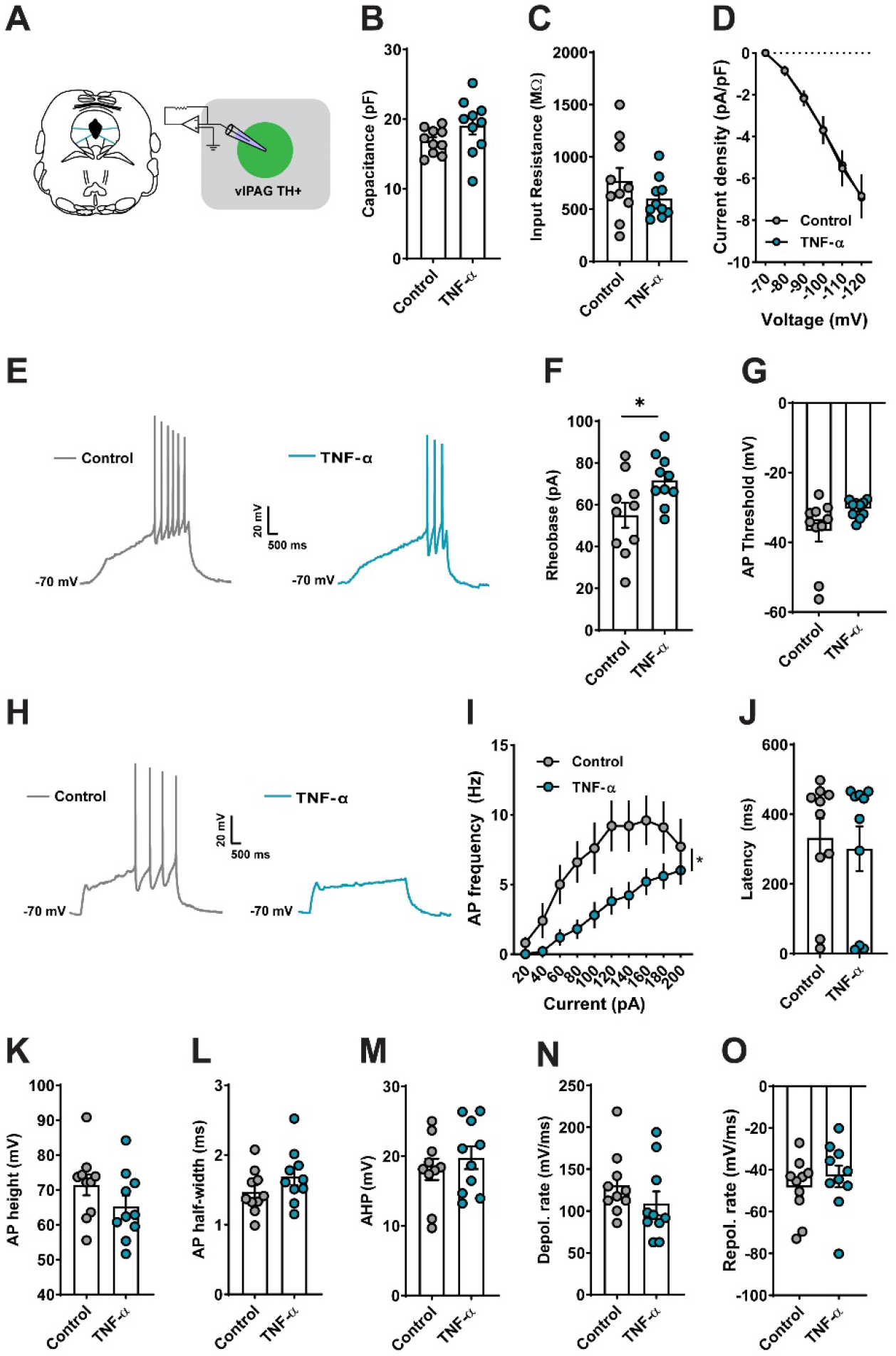
TNF-α incubation decreases excitability of vPAG Dopamine neurons. **(A)** Schematics for whole-cell recordings from vPAG DA (4-6 mice per group) following pre-incubation of slices in TNF-α (100 ng/ml). **(B-D)** There was no change in cell capacitance, input resistance or current-voltage relationship between the two groups. **(E)** Representative traces of vPAG DA neurons in response to a 120 pA/s current ramp while injecting a constant current to hold the cells at -70 mV. **(F)** TNF-α significantly increased the rheobase without altering the AP threshold **(G). (H)** Representative traces of action potentials fired across a range of current steps while injecting a constant current to hold the cells at -70 mV. **(I)** There was a significant interaction between the frequency of action potential fired and TNF-α exposure. Similar to vPAG vGAT neurons, TNF-α did not alter action potential kinetics of vPAG DA neurons (**J-O)**. Data expressed as Mean ± SEM.*p<0.05.

### TNF-α reduces excitatory neurotransmission on vPAG GABA neurons

TNF-α is known to have direct effects on glutamate transmission and has been shown to alter the expression of AMPA receptors on synapses (Beattie et al. 2002; Lewitus et al. 2014; Stellwagen et al. 2005b). Therefore, we evaluated whether TNF-α influences glutamatergic synaptic transmission on both vGAT (Fig. 3) and DA neurons (Fig. 4) in the vPAG. We found that TNF-α decreased spontaneous excitatory synaptic transmission onto vPAG GABA neurons (n=14 cells from 4-5 mice per group). Specifically, we observed significant differences in the cumulative inter-event distribution of sEPSCs, with TNF-α shifting the inter-event distribution to longer intervals (Fig. 3B; p<0.001; Kolmogorov-Smirnov test). There was also a strong trend toward reduction in mean sEPSC frequency in the TNF-α group (Fig. 3C; p=0.0531). Unlike sEPSC frequency, there was no significant effect of TNF-α on the sEPSC amplitude. This can be seen from the lack of impact of TNF-α on the cumulative distribution of amplitude (Fig. 3D; p=0.0689; Kolmogorov-Smirnov test) or the mean amplitude of sEPSCs (Fig. 3E; p=0.0671; unpaired t-test with Welch’s correction). Also, TNF-α significantly decreased the excitatory synaptic drive (calculated as frequency x amplitude) onto vPAG GABA neurons (Fig. 3F; p=0.0285; unpaired t-test with Welch’s correction). Thus, TNF-α incubation leads to a reduction in action-potential dependent release of glutamate onto vPAG GABA neurons.

**Figure 3.**
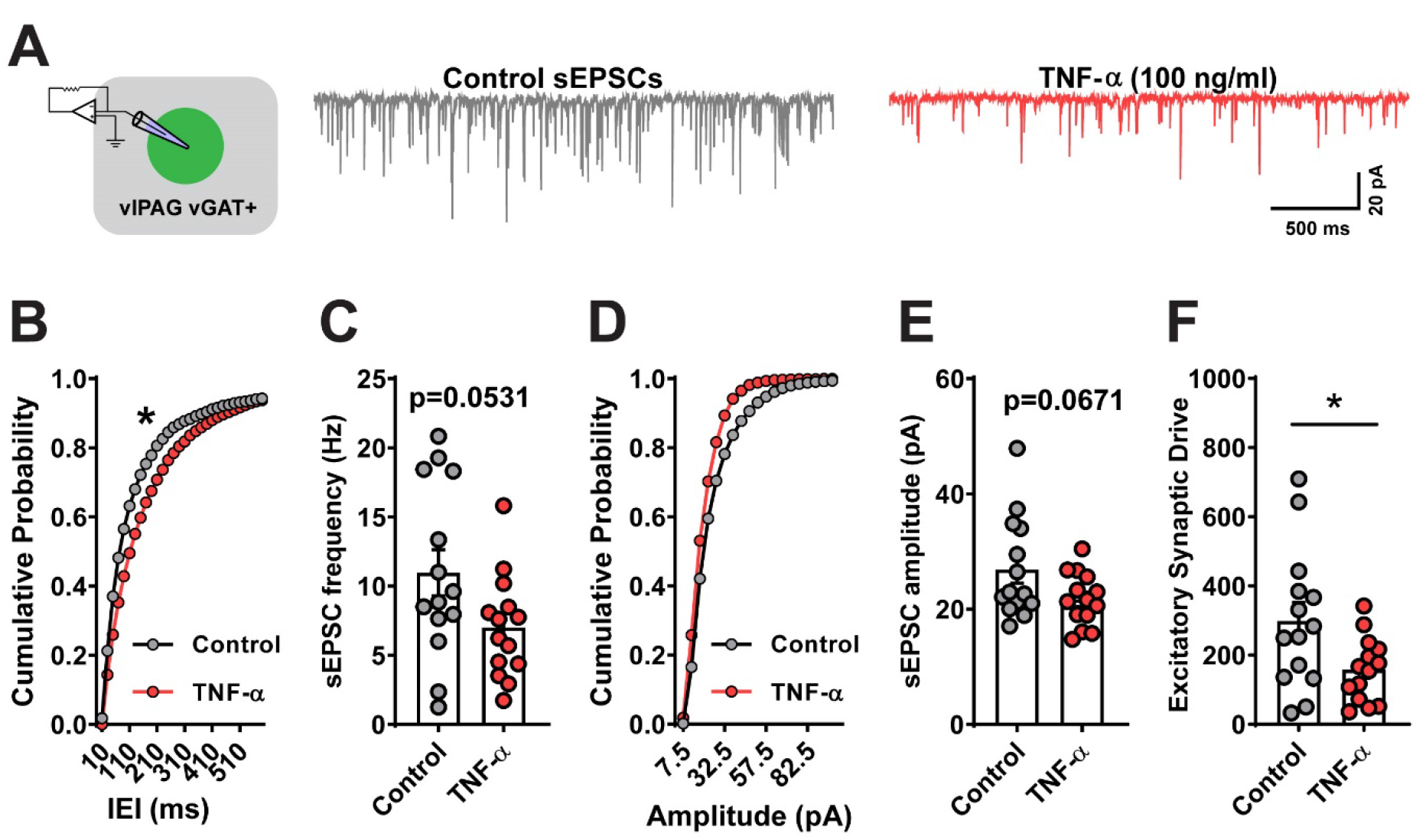
TNF-α incubation reduces spontaneous excitatory synaptic drive on vPAG GABA neurons. **(A)** Representative traces of sEPSCs from vPAG vGAT neurons (n= 14 cells group from 5 mice each) recorded either in ACSF or in the presence of TNF-α (100 ng/ml). A reduction in sEPSC frequency **(C)** was accompanied by a rightward shift in the distribution of sEPSC inter-events interval **(B)** following pre-incubation of slices in TNF-α. **(D-E)** There was a non-significant decrease in sEPSC amplitude onto vPAG vGAT neurons in the presence of TNF-α. **(F)** TNF-α decreased the excitatory synaptic drive onto vPAG GABA neurons. Data expressed as Mean ± SEM.*p<0.05.

**Figure 4.**
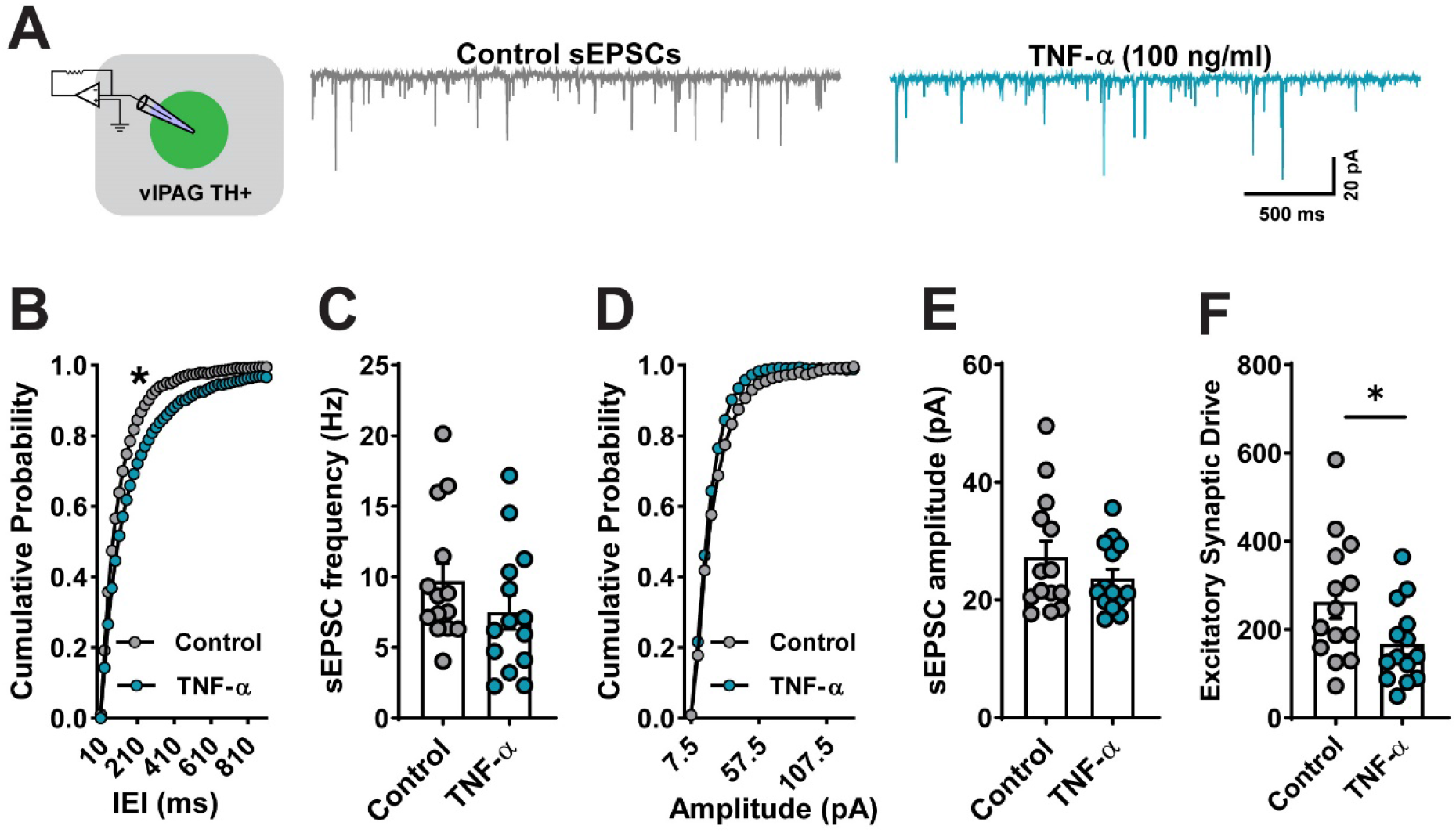
Effects of TNF-α incubation on spontaneous excitatory synaptic transmission in vPAG Dopamine neurons. **(A)** Representative traces of sEPSCs from vPAG DA neurons (n=9-11 cells from 4-5 mice per group) recorded either in ACSF or in the presence of TNF-α (100 ng/ml). TNF-α induced a rightward shift in the distribution of sEPSC inter-events interval **(B)** without a significant decrease in sEPSC frequency **(C)**. No changes were observed in either distribution of sEPSC amplitude **(D)** or mean amplitude **(E)** in vPAG DA neurons. **(F)** TNF-α decreased the excitatory synaptic drive onto vPAG DA neurons. Data expressed as Mean ± SEM.*p<0.05.

### TNF-α has a modest effect on excitatory neurotransmission on vPAG Dopamine neurons

We next examined whether TNF-α alters glutamatergic synaptic transmission onto vPAG DA neurons (Fig. 4; n=14 cells from 5 mice per group). Pre-incubation in TNF-α resulted in a significant shift in the inter-event distribution to longer intervals (Fig. 4B; p= p<0.001; Kolmogorov-Smirnov test) without any changes in mean sEPSC frequency (Fig. 4C; p=0.2167). Additionally, no change was observed in either the distribution (Fig. 4D; p=0.1766; Kolmogorov-Smirnov test) or the mean amplitude of sEPSCs (Fig. 4E; p=0.2462). Like vGAT neurons, TNF-α significantly reduced the excitatory synaptic drive onto vPAG DA neurons (Fig. 4F; p=0.0422). These findings indicate that TNF-α has a modest but significant impact on spontaneous glutamatergic transmission onto vPAG DA neurons.

### TNF-α has no effect on inhibitory neurotransmission on vPAG Dopamine neurons

Evidence from the literature on synaptic connectivity within the PAG suggests that PAG output neurons are tonically inhibited by local GABA neurons (Moreau and Fields 1986; Vaughan et al. 1997). Based on the effects of TNF-α incubation on both excitability and synaptic transmission onto vPAG GABA neurons, we hypothesized that TNF-α would also alter inhibitory drive onto vPAG DA neurons. To investigate this, we isolated spontaneous IPSCs onto DA neurons in the vPAG (Fig. 4A; n=9-12 cells from 3-4 mice per group). Contrary to our hypothesis, TNF-α did not affect inhibitory drive onto DA neurons. Specifically, we observed no differences in either cumulative inter-event distribution of sIPSCs (Fig. 5B; p=0.7937; Kolmogorov-Smirnov test) or mean frequency of sIPSCs (Fig. 5C; p=0.2580). Similarly, TNF-α had no effect on either distribution (Fig. 5C; p=0.1539; Kolmogorov-Smirnov test), the mean sIPSC amplitude (Fig. 5D; p=0.5828) or the inhibitory synaptic drive onto vPAG DA neurons (Fig. 5F; p=0.3325; unpaired t-test with Welch’s correction). Taken together, our results suggest that TNF-α selectively modulates spontaneous glutamate release onto DA neurons without affecting action-potential dependent GABA release.

**Figure 5.**
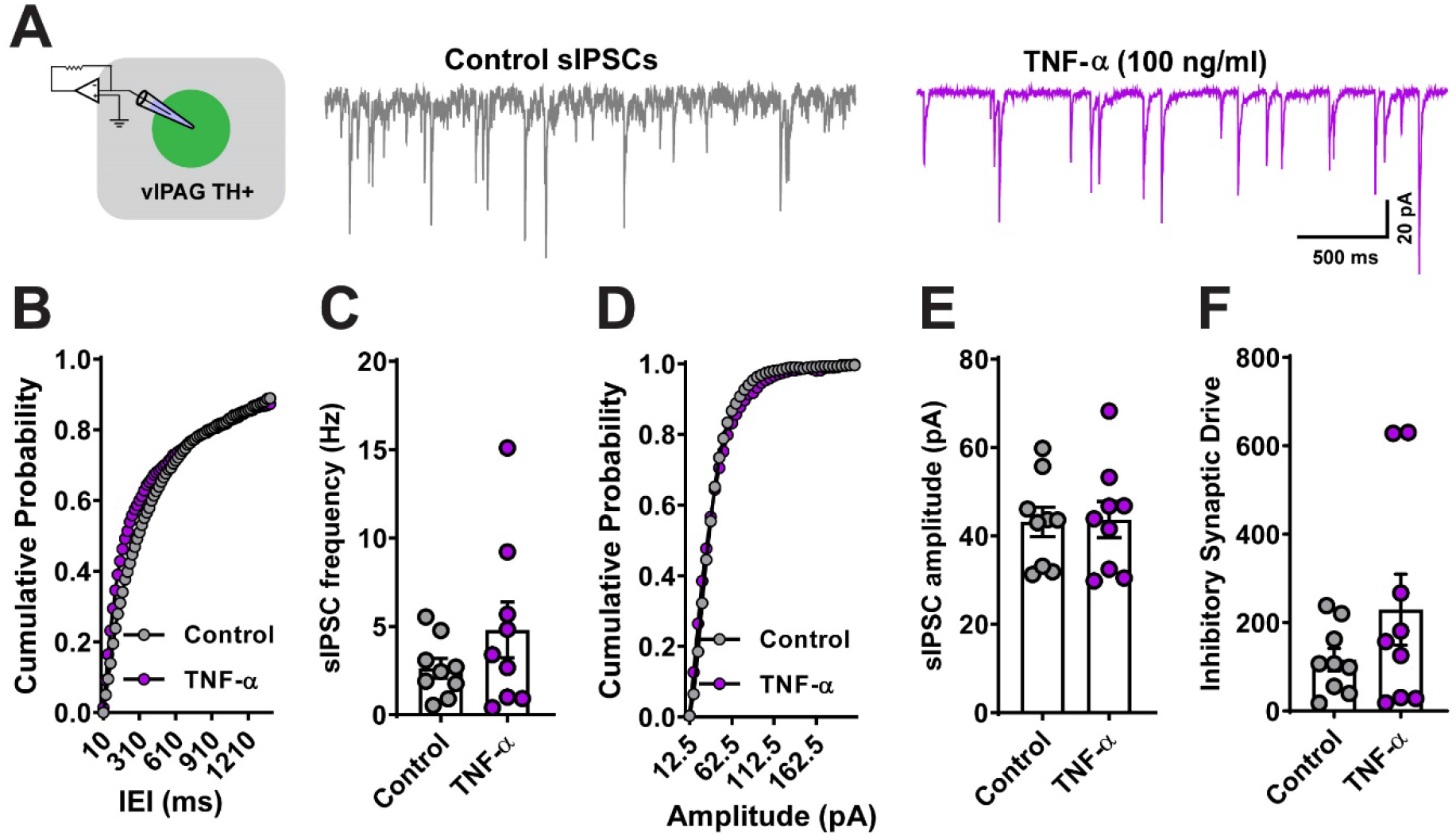
TNF-α incubation does not affect spontaneous inhibitory synaptic transmission in vPAG Dopamine neurons. **(A)** Representative traces of sIPSCs from vPAG DA neurons (n=9 cells from 3 mice per group) recorded either in ACSF or in the presence of TNF-α (100 ng/ml). Cumulative probability plots showing no change in the distribution of sIPSC inter-events interval **(B)** and peak amplitude **(D)**. There were no between-group differences in sIPSC parameters: mean frequency **(C)** and mean amplitude **(E)** in the presence of TNF-α. **(F)** TNF-α did not alter the inhibitory synaptic drive onto vPAG DA neurons. Data expressed as Mean ± SEM.*p<0.05.

## Discussion

The present set of experiments tested the neuromodulatory effects of human recombinant TNF-α on two distinct neuronal populations in the ventral division of PAG. Here we report that incubation of slices in TNF-α altered the intrinsic properties of vPAG GABA neurons resulting in increased excitability but reduced excitatory drive onto these neurons. Conversely, TNF-α significantly decreased the excitability of vPAG DA neurons. Additionally, TNF-α lowered excitatory drive onto vPAG DA neurons but did not impact inhibitory transmission.

TNF-α is a homotrimer protein of 17kDa produced centrally during a variety of inflammatory pathologies and can signal through TNF receptors 1 and 2 (Kinouchi et al. 1991). In addition to its role in various pathologies, it can also modulate neuronal function at low physiological levels (Stellwagen and Malenka 2006). Evidence from the literature suggests that TNF-α is capable of modulating both presynaptic and postsynaptic signaling in neurons. In both rat and mouse hippocampus, TNF-α (across a range of concentrations) increases AMPA/NMDA ratio and glutamate release probability through surface trafficking of AMPA receptors (Beattie et al. 2002; Santello et al. 2011; Stellwagen et al. 2005b). Interestingly, there are known regional differences in TNF-α modulation of neuronal excitability since in the dorsolateral striatum, TNF-α drives the internalization of AMPA receptors and reduces synaptic strength (Lewitus et al. 2014). Similarly, in lateral habenula, TNF-α-mediates a reduction in AMPA/NMDA ratio that drives reduced sociability associated with morphine withdrawal (Valentinova et al. 2019). Here, we demonstrate that even within the same brain region, TNF-α differentially modulates neuronal function in a cell-type-specific manner.

We found incubation of vPAG slices in TNF-α (100ng/ml) resulted in increased rheobase and decreased firing rate of DA neurons while lowering the rheobase of GABA neurons. The mechanisms through which TNF-α alters the intrinsic excitability of these neurons remain unknown. It is possible that TNF-α may influence a nonselective cation channel. Previous studies in dorsal root ganglion (Czeschik et al. 2008), cortical neurons (Chen et al. 2015), and subfornical organ (Simpson and Ferguson 2017) suggest a potential involvement of voltage-gated sodium and calcium channels in driving TNF-mediated neuronal excitability. A recent study in the cerebellum showed TNF receptor 1-mediated downregulation of SK channels as a potential mechanism of increased excitability (Yamamoto et al. 2019). While these studies have identified possible ion channels involved in TNF-α-mediated hyperexcitability, not much is known about the mechanism behind TNF-α driven hypoexcitability. Future studies examining the mechanism behind the hyperpolarization induced by TNF-α may provide insight into the complex role of central TNF-α signaling. Our study also indicates a modest reduction in excitatory synaptic drive onto both GABA and DA neurons. In vPAG GABA neurons, there was a strong trend toward a decrease in both sEPSC frequency and amplitude along with a significant rightward shift in the distribution of sEPSC events. Our finding is consistent with the known role of TNF-α in altering pre and postsynaptic glutamatergic transmission, but it is beyond our scope to address it further.

### Conclusions

These experiments describe the impact of exogenous TNF-α on PAG neuronal activity. Constitutive TNF-α levels in the brain are thought to be low (in the picomolar range) and controlled by local glial cells (Santello et al. 2011). As such, the use of a non-physiological concentration of TNF-α(100 ng/ml) is a limitation of our study and should be taken into consideration while interpreting these findings. However, pathological brain states can induce massive production of proinflammatory cytokines (McCoy and Tansey 2008; Ren et al. 2011; Sud et al. 2007); thus, incubation of slices in saturating levels of TNF-α could potentially reflect plasticity associated with diseased conditions(McCoy and Tansey 2008; Ren et al. 2011; Sud et al. 2007) More in-depth work is needed to demonstrate whether these findings translate under physiological concentrations of TNF-α.

We decided to focus on TNF-α-mediated synaptic alterations in female mice since most studies use male subjects, limiting our understanding of mechanistic variations in females. While it is beyond the scope of the present study, it will be intriguing to determine whether there are sex-related differences in TNF-α modulation of the PAG circuitry. Overall, our findings suggest a potential mechanism whereby pathophysiological concentrations of TNF-α could lead to altered PAG function. Given the known sex differences in pain sensitivity and accumulating evidence pointing toward activated microglia in female PAG as a contributor to sex differences in morphine analgesia, this study provides a mechanistic framework that may provide insight into sex-dependent modulation of pain. As work from our lab and others have demonstrated, these neurons play an opposing role in pain modulation. Either activation of GABA neurons or inhibition of DA neurons results in increased nociception. Therefore, it is plausible that in females, chronic pain results in increased activation of microglia and release of pathophysiological levels of TNF-α, which suppresses the neuronal activity of vPAG DA neurons while increasing the excitability of GABA neurons with the net result being increased nociception and reduced morphine efficacy. Future work is needed to determine the mechanisms by which TNF-α signaling may be upregulated in this region and the behavioral consequences of selective manipulation of PAG-specific TNF-α signaling.

